# Human islet expression levels of Prostaglandin E_2_ synthetic enzymes, but not prostaglandin EP3 receptor, are positively correlated with markers of β-cell function and mass in non-diabetic obesity

**DOI:** 10.1101/2021.02.03.429205

**Authors:** Nathan A. Truchan, Rachel J. Fenske, Harpreet K. Sandhu, Alicia M. Weeks, Chinmai Patibandla, Benjamin Wancewicz, Samantha Pabich, Austin Reuter, Jeffrey M. Harrington, Allison L. Brill, Darby C. Peter, Randall Nall, Michael Daniels, Margaret Punt, Cecilia E. Kaiser, Elizabeth D. Cox, Ying Ge, Dawn B. Davis, Michelle E. Kimple

## Abstract

Elevated islet production of prostaglandin E_2_ (PGE_2_), an arachidonic acid metabolite, and expression of Prostaglandin E_2_ Receptor subtype EP3 (EP3) are well-known contributors to the β-cell dysfunction of type 2 diabetes (T2D). Yet, many of the same pathophysiological conditions exist in obesity, and little is known about how the PGE_2_ production and signaling pathway influences non-diabetic beta-cell function. In this work, plasma arachidonic acid and PGE_2_ metabolite levels were quantified in a cohort of non-diabetic and T2D human subjects to identify their relationship with glycemic control, obesity, and systemic inflammation. In order to link these findings to processes happening at the islet level, cadaveric human islets were subject to gene expression and functional assays. Interleukin-6 (IL-6) and cyclooxygenase-2 (COX-2) mRNA levels, but not those of EP3, positively correlated with donor body mass index (BMI). IL-6 expression also strongly correlated with the expression of COX-2 and other PGE_2_ synthetic pathway genes. Insulin secretion assays using an EP3-specific antagonist confirmed functionally-relevant up-regulation of PGE_2_ production. Yet, islets from obese donors were not dysfunctional, secreting just as much insulin in basal and stimulatory conditions as those from non-obese donors as a percent of content. Islet insulin content, on the other hand, was increased with both donor BMI and islet COX-2 expression, while EP3 expression was unaffected. We conclude up-regulated islet PGE_2_ production may be part of the β-cell adaption response to obesity and insulin resistance that only becomes dysfunctional when both ligand and receptor are highly expressed in T2D.

---

Fundamentally, type 2 diabetes mellitus (T2D) results from a failure of the insulin-secreting pancreatic β-cells to compensate for peripheral insulin resistance. Obesity is the most common co-morbidity found in individuals with insulin resistance. While there exists debate about whether one condition precedes the other, the physiological changes that often occur in the obese, insulin-resistant state (e.g., systemic inflammation, dyslipidemia, hyperinsulinemia, mild hyperglycemia) simultaneously induce β-cell stress and increase β-cell workload, forcing the β-cell to initiate a compensatory response in order to function, replicate, and survive^*1–3*^. Whether the β-cell can initiate and continue this adaptive program is the key determinant of the progression to T2D^*2*^.

Cyclic adenosine monophosphate (cAMP) is a well-characterized potentiator of glucose-stimulated insulin secretion (GSIS) and promotes a number of proliferative and survival pathways in the β-cell^*4–8*^. Numerous changes in cAMP homeostasis occur in a highly compensating β-cell, all with the central theme of increasing cAMP production, decreasing cAMP degradation, or promoting signaling through downstream effectors^*7,9–12*^. Our and others’ previous studies suggest that signaling through Prostaglandin EP3 receptor (EP3), a G protein-coupled receptor for the arachidonic acid (AA) metabolite, prostaglandin E_2_ (PGE_2_), is up-regulated by the pathophysiological conditions of T2D, negatively influencing β-cell function and mass through its inhibition of adenylyl cyclase^*13–21*^. Islets isolated from T2D mice and human organ donors express more EP3 and produce more PGE_2_ than islets isolated from non-diabetic controls^*13, 18*^, and treating these islets with an EP3 receptor antagonist, L798,106, potentiates GSIS and restores their secretory response to glucose and glucagon-like peptide 1 receptor (GLP1R) agonists^*13*^. As GPCRs form the largest class of druggable targets in the human genome, we and others have previously proposed EP3 as a putative therapeutic target for the β-cell dysfunction of T2D^*9, 13, 14, 18, 22–25*^. Yet, other studies have found no correlation of EP3 with human organ donor obesity and/or T2D status, and PGE_2_ has other downstream effects shown to be important in β-cell replication and/or survival^*9, 14*^. Finally, while the effects of EP3 signaling in metabolically healthy islets and T2D islets have been well-characterized, little is known on the role of PGE_2_ production and signaling in the β-cell highly compensating for obesity and insulin resistance. As many of the same metabolic derangements of T2D are present in severe insulin resistance (e.g., hyperglycemia, dyslipidemia, systemic inflammation, etc.), this is a critical issue to address.

To our knowledge, a comprehensive analysis of the PGE_2_ production and EP3 signaling pathways and their correlations with β-cell function and mass have never been completed in pancreatic islets of non-diabetic human donors spanning the obesity spectrum. In this work, we profiled the islet mRNA expression of genes encoding EP3 and PGE_2_ production pathway enzymes using a comprehensive set of islets obtained from 40 non-diabetic human organ donors distributed across a BMI range of ~22-45 kg/m^2^. Just over half of these islet preparations were also used in glucose-stimulated and incretin-potentiated insulin secretion assays to determine the impact of endogenous EP3 signaling on β-cell function with insulin content serving as a surrogate of β-cell mass. The relationship(s) between plasma PGE_2_ metabolite levels, glycemia, obesity, and systemic inflammation in a clinical cohort of non-diabetic and T2D subjects spanning a similar BMI spectrum supports the clinical relevance of our findings.

## Results

### Plasma PGE_2_ metabolite (PGEM) levels correlate strongly with T2D disease status, but are also elevated in obese non-diabetic subjects in a clinical cohort

Biobanked plasma samples from 19 conservatively-treated T2D subjects with well-controlled disease and 16 non-diabetic (ND) controls, well-matched for age and BMI, were selected for downstream analysis. Approximately equal numbers of male and female subjects were represented between groups, of similar age mean age and age range, with the BMI range of both groups spanning normal weight (BMI < 24.9) to high-risk obesity (BMI ≥ 40) (**Table 1**).

**Table 1:** Demographics of Non-diabetic (ND) and T2D study subjects for plasma metabolomics analyses.

|  |  |  |  |  |
| --- | --- | --- | --- | --- |
| **Cohort Demographics** | **All** | **Non-Diabetic (ND)** | **T2D** | **P-value** |
| **Subjects (N)** | 35 | 16 | 19 | - |
| **Male, N (%)** | 85 (51%) | 5 (31%) | 8 (42%) | - |
| **Age (y), mean ± SD (range)** | 54.1 ± 10.5 (29-73) | 49.8 ± 10.8 (31-70) | 55.6 ± 8.1 (41-67) | 0.0813 |
| **BMI (kg/m<sup>2</sup>), mean ± SD (range)** | 31.6 ± 8.3 (19.5-61.9) | 31.6 ± 5.3 (24.7-42.2) | 34.3 ± 6.2 (23.4-47.3) | 0.1738 |
| **HgA1c (%), mean ± SD (range)** | - | 5.3 ± 0.3 (4.8-5.8) | 6.7 ± 0.7 (5.8-8.1) | <0.0001 |

PGE_2_ in clinical biosamples undergoes rapid metabolism to form 13,14-dihydro-15-keto prostaglandin E_2_, which undergoes a variable degree of degradation to form 13,14-dihyrdro-15-keto prostaglandin A_2_ depending on the type of biofluid, processing, and storage conditions. The Prostaglandin E Metabolite (PGEM) ELISA (Cayman Chemical Company) converts both of these metabolites to a single, stable derivative correlating with initial PGE_2_ levels. Mean plasma PGEM was significantly elevated in the T2D group vs. ND group (an approximate 200% increase) (**Figure 1A**). Obesity status had no additive effect on PGEM levels in T2D subjects, but in ND subjects, mean plasma PGEM was approximately 40% higher in obese subjects vs. non-obese (**Figure 1A**).

**Figure 1.**
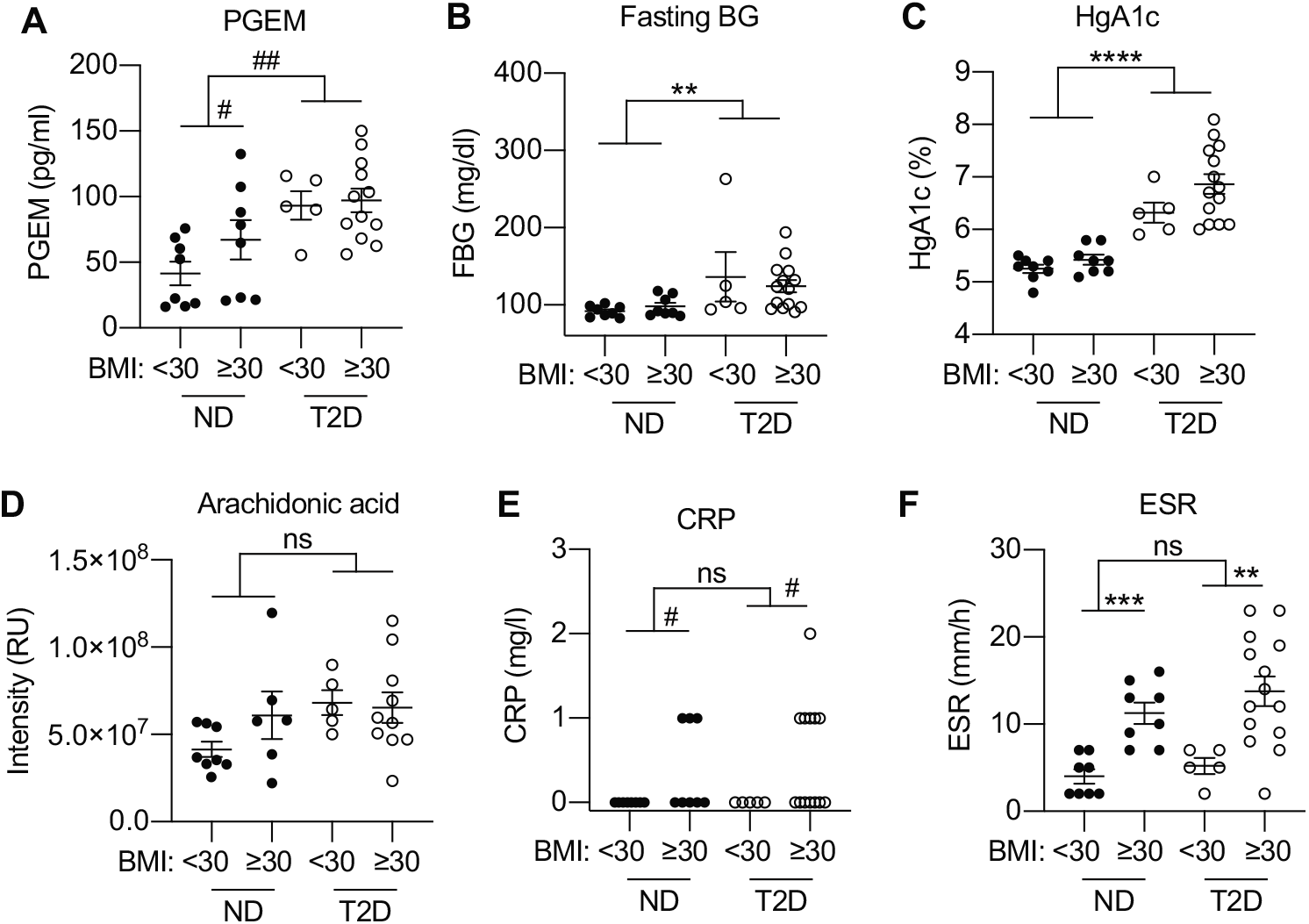
Plasma PGE_2_ metabolite (PGEM) levels are independently associated with obesity and T2D status in a clinical cohort. Fasting plasma samples were collected from 19 subjects with T2D well-controlled with diet and lifestyle changes or metformin monotherapy and 16 nondiabetic (ND) controls. A, Plasma PGEM levels. B, Fasting blood glucose. C, HgA1c. D, Plasma arachidonic acid, E, Serum C-reactive protein (CRP). F, Blood erythrocyte sedimentation rate (ESR). Unless otherwise indicated, data between ND and T2D groups, and, within each group, non-obese and obese donors was compared by two-tail T-test, except when data were not normally distributed (indicated by #). If not shown, a comparison was not statistically significant. **, p < 0.01; ***, p < 0.001, and ****, p < 0.0001. ns = not significant.

Expression of the rate-limiting enzymes in PGE2 production, cyclooxygenase (COX) 1 and 2, are induced in hyperglycemic conditions^*17, 18, 26, 27*^. As expected, fasting blood glucose (FBG) and HgA1c were significantly higher in T2D vs. ND subjects, but obesity had no significant effect in either group (**Figure 1B,C**). Therefore, any increase in plasma PGEM levels in obese ND subjects was not mediated by elevated glucose levels, whether acute or chronic.

AA cleaved from plasma membrane phospholipids by phospholipase A2 (PLA2) is the substrate for COX enzymes. AA is an essential fatty acid obtained from the diet. Plasma AA levels were quantified using an ultrahigh resolution flow-injection ionospray (FIE) Fourier transform ion cyclotron resonance (FTICR) mass spectrometry (MS)-based platform developed and validated previously for T2D biosamples^*17, 28*^. While the mean AA levels in non-obese ND subjects were the lowest of all four groups, plasma AA levels were relatively unaffected by either obesity or T2D status (**Figure 1D**).

Besides high glucose, proinflammatory cytokines are another well-known inducer of COX expression and activity^*29–33*^. Two clinical tests that measure a patient’s inflammatory state were used in our clinical study: C-reactive protein (CRP) and erythrocyte sedimentation rate (ESR). CRP is a non-specific indicator of inflammation levels and is released from the liver in response to acute and chronic injury and/or inflammation. In our cohort, CRP levels were not significantly different between ND and T2D subjects, but were elevated in both groups with obesity, with only obese subjects having CRP levels > 0 (**Fig. 1E**). ESR is a non-specific test of systemic inflammation and was strongly elevated in obese subjects, whether ND or T2D (**Figure 1F**).

### Relative mRNA levels of key enzymes in the PGE_2_ production pathway are positively correlated with non-diabetic donor BMI, obesity status, and islet *IL6* expression

The pro-inflammatory cytokine, interleukin-6 (IL-6), is increased in obesity and associated with β-cell compensation^*34–36*^. Relative quantitative PCR (qPCR) for IL-6, EP3, and PGE_2_ synthetic pathway genes was performed on cDNA samples from a panel of islet preparations from nondiabetic human organ donors spanning a BMI range of 22.8-44.7: nearly identical to that in our clinical cohort. The set included 27 male and 13 female donors, with a mean age of 42.6 years and mean BMI of 31.5 kg/m^2^ (**Table 2**). The difference between the mean ages of non-obese (40.7 years) and obese donors (44.0 years) was not statistically different. Seventeen islet preparations were from donors with BMI < 30. Five were from lean donors (BMI < 25), with no donors being underweight (BMI < 18.5), and 12 were from overweight donors (BMI 25-29.9). Of the 23 obese islet donors, 13 were in the low-risk obesity category (BMI < 35), 8 were in the moderate-risk category (BMI 35 – 39.9), and 2 were classified as high-risk obesity (BMI ≥ 40). PCR primer sequences can be found in **Table S1** and islet donor and isolation parameters can be found in **Table S2**).

**Table 2:**
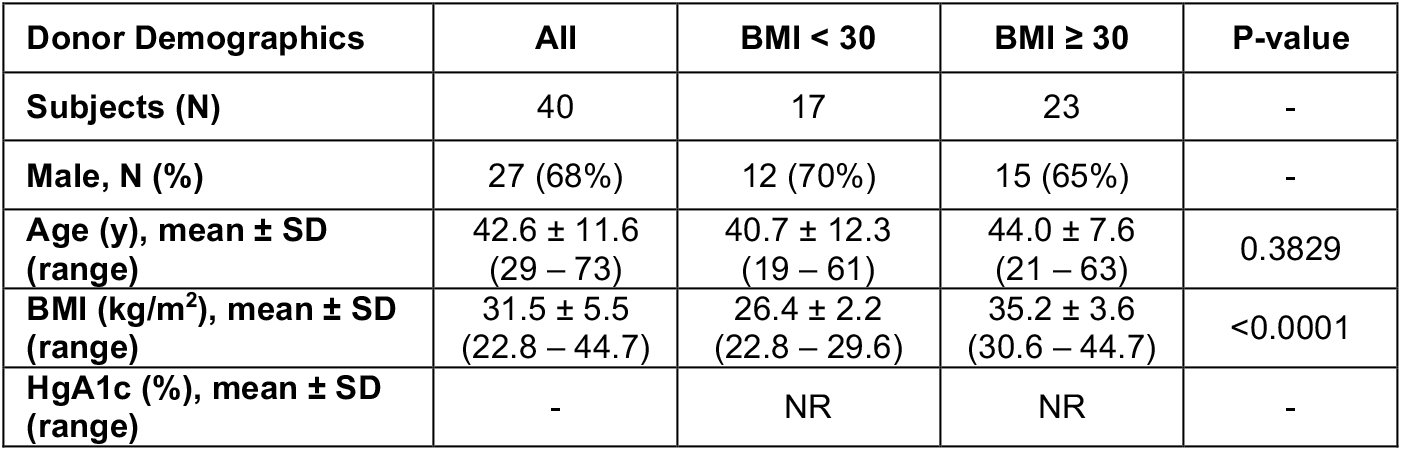
Donor demographics of islet preparations used in gene expression and insulin secretion assays (i.e., Islet Set 1). NR, not reported.

The relationship between islet gene expression and BMI was determined using two methods: (1) linear regression and (2) binning samples by donor obesity status (BMI < 30 and BMI > 30; N=17 non-obese and N=23 obese donors in each set) and performing a two-tail T-test. The complete results of this analysis can be found in **Table S3**. Besides IL-6 (*IL6*) and COX-2 (*PTGS2*) (**Figure 2A,B**), none of the other genes were significantly correlated with BMI, including EP3 (*PTGER3*) (**Figure 2C**). Because the latter finding is partially in conflict with previously-published work^*13, 14*^, *PTGER3* expression was measured from an independent set of non-diabetic human donor islets with almost identical sex distribution, age range, and BMI range (**Table 4 and Table S4**), with nearly identical results (**Figure 2D**). Finally, *PTGS2* expression was highly correlated with *IL6*, as were all of the other PGE_2_ production genes, save *PTGES3* (**Figure 3**). No relationship was observed between *IL6* and *PTGER3* expression (**Table S2**).

**Figure 2.**
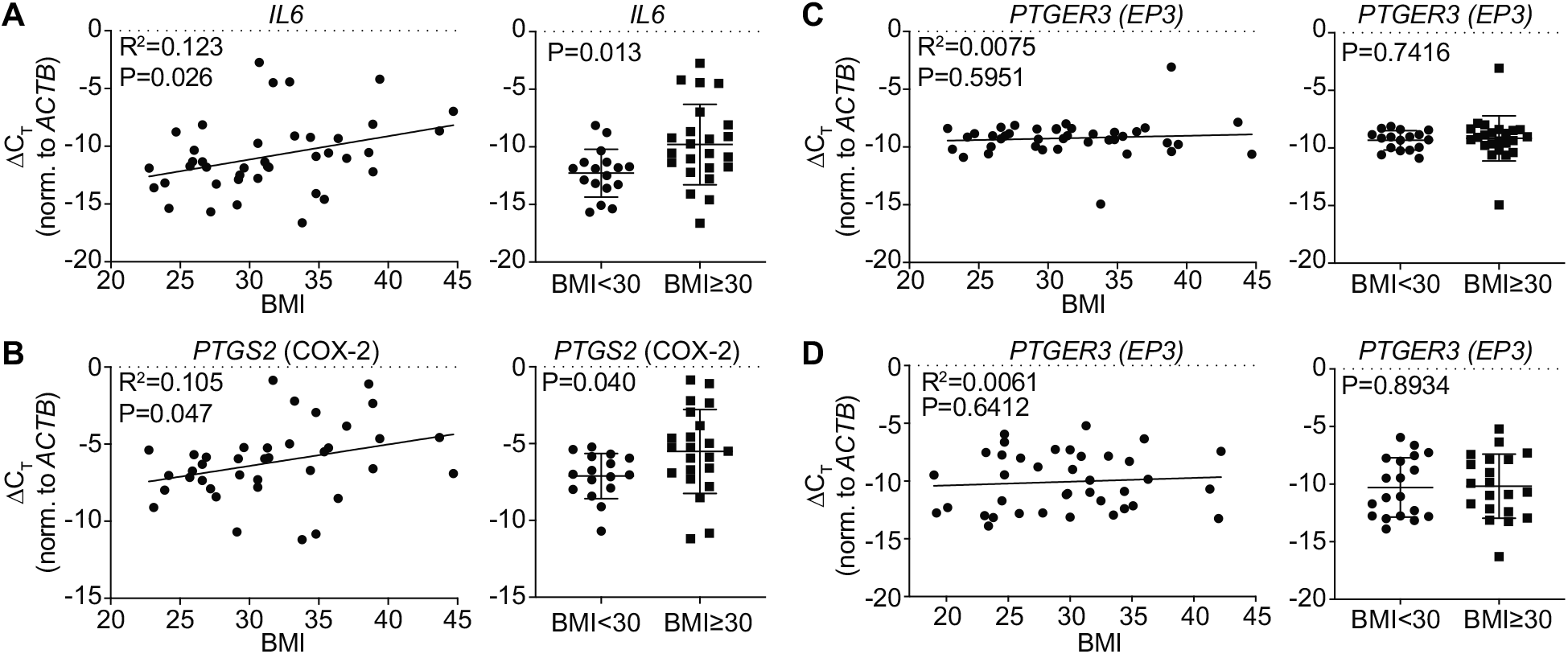
Islet *IL6* and *PTGS2* expression, but not that of *PTGER3*, positively correlates with donor BMI. A-C: Quantitative PCR results from Islet Set 1 for (A) *IL6*, (B) *PTGS2*, and (C) *PTGER3*. D: Quantitative PCR results from Islet Set 2 for *PTGER3*. Data are represented as cycle time normalized to that of β-actin (*BACT*) (ΔC_T_). Data were analyzed by linear regression vs. donor BMI (left panels) or two-tailed t-test by donor obesity status (right panels). The Goodness-of-fit (R^2^) and P-value after linear regression analyses and P-value after twotailed t-test are indicated.

**Figure 3.**
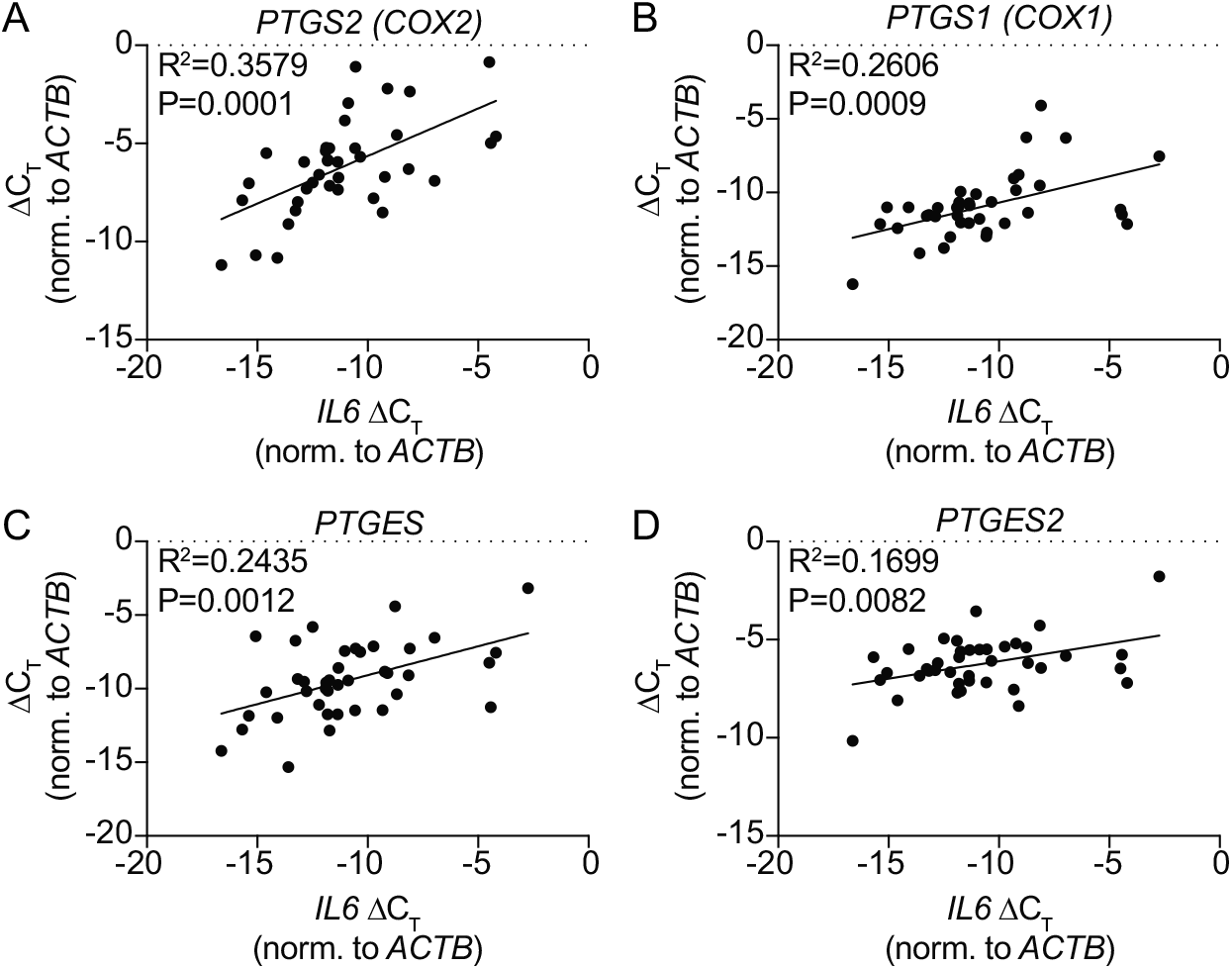
PGE_2_ production genes are positively correlated with *IL6* expression. A-D: Quantitative PCR results from Islet Set 1 for (A) *PTGS2*, (B) *PTGS1*, (C) *PTGES*, and (D) *PTGES2*. Data are represented as cycle time normalized to that of β-actin (*BACT*) (ΔC_T_) and were analyzed by linear regression. The Goodness-of-fit (R^2^) and P-value are indicated.

### An EP3 antagonist promotes glucose-stimulated and incretin-potentiated insulin secretion in islets from obese donors only

Activated EP3 acts as a non-competitive antagonist of GLP1R, blocking its maximal potentiating effect on insulin secretion^*13*^. In order to test the impact of endogenous EP3 signaling on human β-cell function in non-diabetic obesity, we performed glucose-stimulated insulin secretion (GSIS) assays with and without the addition of the GLP1R agonist, exendin-4 (Ex4) and the EP3 antagonist, L798,106. Islets from non-obese and obese donors were both glucose responsive, with significantly more insulin secreted as a percent of content in stimulatory glucose (16.7 mM) vs. basal glucose (1.7 mM) (**Figure 4**). Comparing the results from non-obese to obese donor islets, GSIS was higher in every condition tested, with GSIS in 1.7 mM glucose and 16.7 mM glucose plus L798,106 being significantly so. Finally, only in islets from obese donors did L798,106 and Ex4 treatment synergize to enhance GSIS above 16.7 mM glucose alone. These results confirm the elevated expression of PGE_2_ synthetic genes in non-diabetic obesity is of functional consequence, although islets from the obese donors in the cohort used in our studies were not dysfunctional.

**Figure 4.**
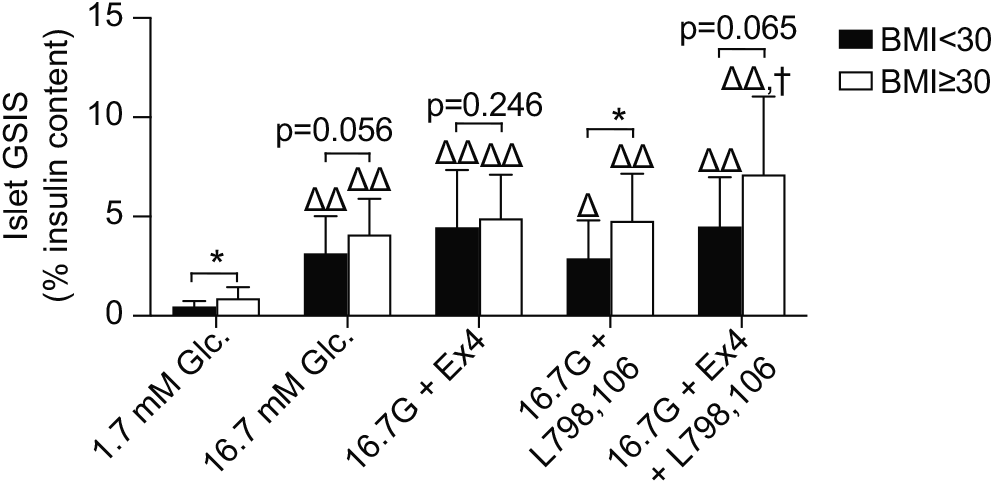
An EP3 antagonist promotes glucose-stimulated and incretin-potentiated insulin secretion in islets from obese donors only. Acute islet insulin secretion by donor obesity status (N=11, BMI < 30 and N=10, BMI ≥ 30) in the presence of 1.7 mM glucose, 16.7 mM glucose, or 16.7 mM glucose with 10 nM Ex4, 20 μM L798,106, or both. Data are represented as % insulin secreted as normalized to content and plotted as mean ± SD. Within treatments, BMI groups were compared by twoway t-test. Within BMI groups, data were compared by one-way paired ANOVA with Holm-Sidak test post-hoc to correct for multiple comparisons. *, p<0.05 for BMI < 30 vs. BMI ≥ 30. Δ, P<0.05 and ΔΔ, P<0.01 as compared to 1.7 mM glucose. †, p<0.05 as compared to 16.7 mM glucose. Unless indicated, a comparison was not statistically significant.

### Islet insulin content is positively correlated with donor *BMI* and *PTGS2* expression

The GSIS assays in Figure 5 were performed on single islets adhered to the vertex of a 96-well V-bottom tissue-culture treated microtiter plate, and each islet’s insulin secretion was normalized to its own insulin content. This method has been previously optimized by our laboratory to produce high-quality, reproducible results from mouse and human islets^*37*^. With the standard curve, this provides up to 76 biological replicates per islet preparation to calculate mean islet insulin content. Insulin content itself is significantly correlated with BMI, both linearly (p=0.0018; R^2^=0.408) and by obesity status (p<0.01) (**Figure 5A,B**). Notably, insulin content was also significantly correlated with *PTGS2* expression (P=0.0005; R^2^=0.4766). None of the other genes had a significant correlation with islet insulin content, although *IL6* was the second-most highly correlated after *PTGS2* (p=0.172; R^2^=0.096) (**Table S2**).

**Figure 5.**
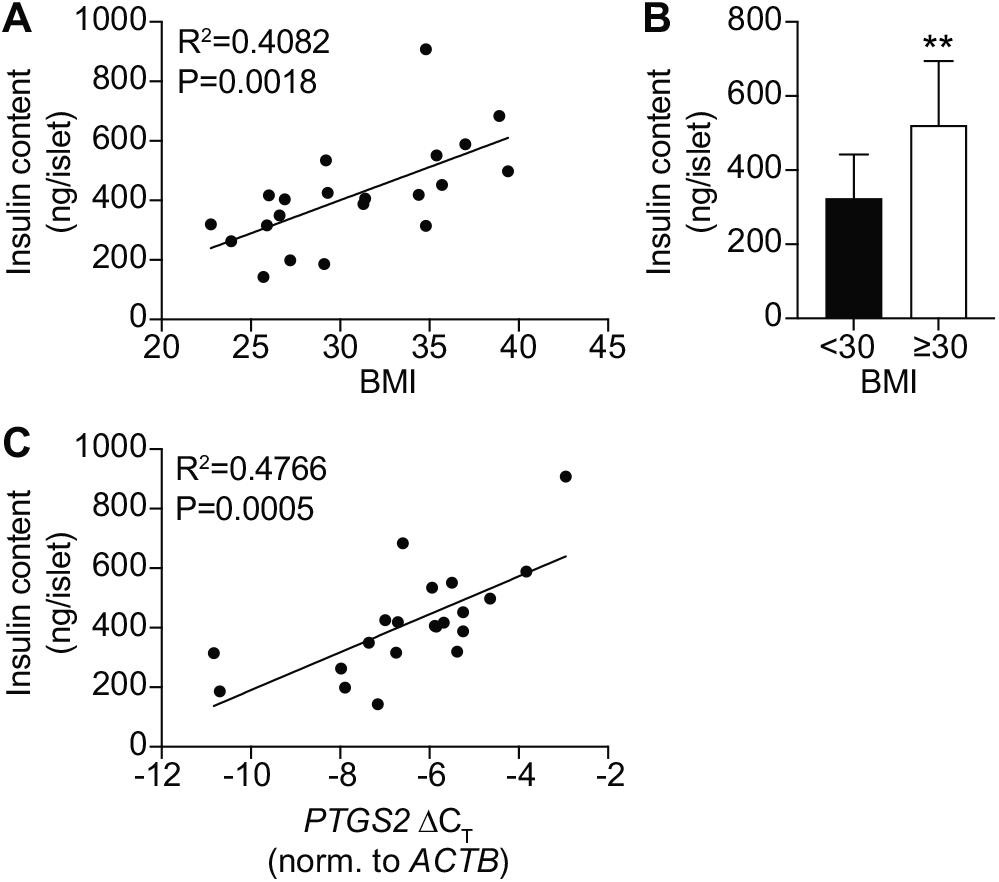
Islet insulin content is positively correlated with BMI and *PTGS2* expression. A & B, Islet insulin content (ng/islet) was subject to linear curve-fit analysis vs. donor BMI (A) or twotailed t-test by donor obesity status (B). C, Islet insulin content was subject to linear curve-fit analysis vs. *PTGS2* expression. In A and C, the goodness-of-fit (R^2^) and p-value for deviation from zero of each of the analyses are indicated. In (B), N=11, BMI < 30 and N=10, BMI ≥ 30. Data represent mean ± SD and were compared by twoway t-test. N=10-11 per group. **, p < 0.01

## Discussion

Insulin resistance, often found with obesity, necessitates the hyperproduction and processing of proinsulin into mature insulin in order to augment insulin secretory capacity, causing significant mitochondrial oxidative and ER stress^*3, 38, 39*^. Together with hormonal dysregulation, dyslipidemia, and systemic inflammation, which also impact the β-cell, downstream adaptive processes need to unfold in order to prevent progression to T2D^*2*^. Many of the factors linked with β-cell adaptation in the obese and/or insulin resistant state(s) have also been proposed as contributing to the development and/or pathophysiology of T2D itself, including our and others’ previously published work on PGE_2_ production and EP3 signaling^*13, 14, 16–22, 24, 40*^.

Elevated circulating levels of PGEM were first linked with diabetes pathophysiology in a study of subjects with type 1 diabetes undergoing diabetic ketoacidosis (DKA)^*41*^. In our small clinical cohort, we found significantly elevated plasma PGEM levels in T2D subjects as compared to ND, but no association of PGEM with severity of hyperglycemia, suggesting the contribution of factors beyond blood glucose levels in the elevated PGEM in T1D DKA. Further, within the ND cohort, but not T2D, we found PGEM associated with BMI. These results both support and contradict previously published works, also in small cohorts. Elisia and colleagues found plasma PGE_2_ levels, as measured by PGE_2_ ELISA, were significantly elevated in samples from 42 nondiabetic obese individuals with BMI > 35 as compared to 35 lean subjects (BMI 18.5 – 24.9)^*42*^. Caveats to these results include a lack of overweight (BMI 25-29.9) subjects being represented, as well as the rapid metabolism of PGE_2_, which typically necessitates PGEM assay used in the current work. Tans and colleagues applied LC/MS to plasma samples from nine lean, ten obese nondiabetic, and 11 T2D subjects, finding no difference in mean PGEM levels between lean and obese non-diabetic subjects vs. a significant elevation in T2D subjects, consistent with our findings^*43*^. Xia and colleagues developed a quantitative LC/MS assay for a panel of polyunsaturated fatty acid-derived eicosanoids, finding plasma levels of a subset of omega-6-derived eicosanoids, including PGE_2_, were highly predictive of T2D diagnosis in a cohort of 20 non-diabetic controls and 20 T2D subjects^*44*^. Within the T2D group, but not the healthy control group, these eicosanoids were associated with BMI and other clinically-relevant T2D parameters, in contrast to our own findings. Finally, Pawelzik and colleagues used an LC/MS approach to analyze urine from 45 obese subjects for 9,15-dioxo-11α-hydroxy-13,14-dihydro-2,3,4,5-tetranor-prostan-1,20-dioic acid (tetranor-PGEM) levels, finding them unrelated to BMI. Tetranor-PGEM did correlate with a higher waist-hip ratio and mildly-elevated HbA1c and 60-min blood glucose levels after glucose challenge, suggestive of pre-diabetes^*45*^. One major caveat to this study, though, is no non-obese subjects were included in the analysis^*45*^.

PGE_2_ is the most abundant natural ligand for the EP3 receptor (encoded by the *PTGER3* gene). COX-1 and COX-2 (officially known as prostaglandin-endoperoxidase 1 and 2: *PTGS1* and *PTGS2*), catalyze the rate-limiting step in the production of PGE_2_ derived from arachidonic acid (AA) incorporated in plasma membrane phospholipids. High glucose, free fatty acids, and/or pro-inflammatory cytokines have all been shown to upregulate the expression and/or activity of enzymes involved in the PGE_2_ synthetic pathway, including phospholipase A2 (PLA2: which cleaves AA from membrane phospholipids), COX-1 and COX-2 (which sequentially convert arachidonic acid to the intermediates PGG_2_ and PGH_2_), and *PTGES*, *PTGES2*, and *PTGES3*, which convert PGH_2_ to PGE_2_^*13, 14, 17–21, 24, 33, 46–49*^. Using BMI as a marker of obesity, we found a positive correlation with *PTGS2* expression, consistent with previously-published results^*13*^. More surprising was the lack of correlation of EP3 gene expression with BMI. We previously published or contributed to two studies characterizing *PTGER3* expression in human islet cDNA samples from non-diabetic human organ donors of varying BMI with seemingly disparate results^*13, 14*^. In Kimple and colleagues, when islets were binned by obesity status, *PTGER3* expression was significantly correlated with donor BMI^*13*^. In the current study, *PTGER3* expression was not correlated with the BMI of the donor, whether linearly or by binning islets by obesity status. This finding was replicated in a separate cohort of 40 non-diabetic human donor islets, and is also consistent with the findings of Carboneau and colleagues^*14*^. As the mean difference in *PTGER3* expression by obesity status in Kimple and colleagues was relatively small (fold-change ≈ 2.8), it is likely this change is of limited biological relevance even if statistically significant.

Pro-inflammatory cytokines have long been understood as contributing to β-cell death and decreased viability^*19–21, 23, 24, 46, 50*^. IL-1β is up-regulated in islets from T2D mice and humans as compared to non-diabetic controls, and treating islets with IL-1β ex vivo significantly up-regulates PGE_2_ production, EP3 expression, or both^*18–21, 33*^. On the other hand, recent reports suggest IL-6 as a multi-potent obesity-associated cytokine that stimulates β-cell adaptation. IL-6 promotes β-cell survival, reduce oxidative stress, and promote glucagon-like peptide 1 (GLP-1)-potentiated insulin secretion^*51–54*^. IL-6 is required for the adaptive proliferative response to high fat diet-induced glucolipotoxicity by paracrine effects on the alpha-cell^*55*^. IL-6 also promotes β-cell autophagy—a survival mechanism working in concert with the adaptive unfolded protein response—through stimulation of alpha-cell GLP-1 production^*53*^. In the current work, we chose islet IL-6 expression as a physiologically-relevant indication of adiposity and the need to induce beta-cell compensation. Our results, although correlative, suggest the possibility of a mechanistic link between islet IL-6 expression and PGE_2_ production. In support of this concept, IL-6 expression has been specifically linked with PGE_2_ produced by COX-2 and not COX-1^*56*^. COX-2 is of particular relevance to the β-cell, as in contrast to other cell types, β-cell COX-2 expression is constitutive^*49*^ and is the major isoform responsible for both endogenous and stimulated β-cell PGE_2_ production^*26, 47, 49, 57*^.

Previous studies, including our own, have shown little-to-no effect of EP3 agonists or antagonists on human islet β-cell function, proliferation, or survival, unless the islets were cultured in glucolipotoxic conditions^*13, 14, 40*^. The comprehensive gene expression and functional assays in our current work confirm up-regulation of PGE_2_ production enzymes in non-diabetic obesity is of functional consequence. Because L798,106 is an EP3-specific competitive antagonist, it has no biological activity of its own: it simply displaces the native agonist. Yet, islets from obese donors were not dysfunctional. In fact, they secreted at least as much insulin as a percent of content as islets from non-obese donors, in addition to having significantly elevated islet insulin content. Increased fractional basal insulin secretion in larger islets or islets from higher BMI organ donors is consistent with a β-cell compensation response to obesity and insulin resistance, which has been observed previously^*58, 59*^. What we did not predict was the strong relationship of islet *PTGS2* expression with preserved beta-cell function and increased insulin content in non-diabetic donors by BMI. Our findings suggest a dose-dependent effect of PGE_2_ production and signaling in obesity and T2D which, if true, has implications in studies of EP3 as a therapeutic target for the prevention and treatment of T2D.

## Materials and Methods

### Materials and Reagents

Sodium chloride (S9888), potassium chloride (P3911), magnesium sulfate heptahydrate (M9397), potassium phosphate monobasic (P0662), sodium bicarbonate (S6014), HEPES (H3375), calcium chloride dehydrate (C3881), exendin-4 (E7144) and radioimmunoassay (RIA)-grade bovine serum albumin (A7888) were purchased from Sigma Aldrich (St. Louis, MO, USA). Anti-insulin antibodies (Insulin + Proinsulin Antibody, 10R-I136a; Insulin + Proinsulin Antibody, biotinylated, 61E-I136bBT) were from Fitzgerald Industries (Acton, MA, USA). The 10 ng/ml insulin standard (8013-K) and assay buffer (AB-PHK) were from MilliporeSigma (Burlington, MA, USA). RPMI 1640 medium (11879–020: no glucose), penicillin/streptomycin (15070–063), and fetal bovine serum (12306C: qualified, heat inactivated, USDA-approved regions) were from Life Technologies (Carlsbad, CA, USA). Dextrose (D14–500) was from Fisher Scientific (Waltham, MA, USA). The RNeasy Mini Kit and RNase-free DNase set were from Qiagen (Valencia, CA, USA). The High-Capacity cDNA Reverse Transcription Kit was from Applied Biosystems (Foster City, CA, USA). FastStart Universal SYBR Green Master mix was from Roche (Indianapolis, IN, USA).

### Human islet preparations

Cultured human islets were obtained from the Integrated Islet Distribution Program (IIDP) under an approved exemption (UW 2012–0865) for non-human subjects research by the UW Health Sciences IRB. One set of 40 islet preparations was used for gene expression and insulin secretion assays, while a second set of 40 was used for confirmation of gene expression only (Set 2 included three flash-frozen islet samples purchased from Beta-Pro LLC, Charlottesville, VA, USA). The unique identifiers for each islet preparation, along with age, sex, BMI, HbA1c, and islet isolation center are listed in Supplementary Tables 1 and 2. All but two islet preparations were associated with a donor HbA1c value. Islet samples for quantitative PCR analysis were handpicked and pelleted on the day of arrival and flash-frozen prior to RNA isolation. Prior to *ex vivo* insulin secretion assays, human islets were cultured for at 24 h in human islet medium (RPMI 1640 medium containing 8.4 mM glucose, supplemented with 10% heat inactivated FBS and 1% penicillin/streptomycin).

### Quantitative PCR assays

150-200 islets from each human islet preparation were washed with PBS and used to generate RNA samples via Qiagen RNeasy Mini Kit according to the manufacturer’s protocol. Copy DNA (cDNA) was generated and relative qPCR performed via SYBR Green assay using primers validated to provide linear results upon increasing concentrations of cDNA template, as previously described^*35*^. The primer sequences used in the qPCR experiments are listed in Table 2.

### Ex vivo islet insulin secretion assays

The day before assay, islets were hand-picked into 96-well V-bottom tissue culture plates and incubated overnight in RPMI growth medium to adhere islets to the bottom of the plate and single-islet GSIS assays performed with and without the addition of the indicated compounds to the stimulation medium, as previously described^*37*^. The secretion medium was collected for insulin secretion assessment and islets were lysed in RLT buffer to measure the insulin content using inhouse-generated insulin sandwich ELISA as described previously^*37*^. In general, secretion medium was diluted 1:20 and content medium diluted 1:200 in order for readings in the linear range of the assay.

### Clinical Study Participant Recruitment and Sample Collection

For the current work, plasma samples from 16 non-diabetic subjects and 19 T2D subjects, the latter treated with diet/lifestyle modifications or metformin monotherapy only, were selected from existing biobanked samples (details follow) for downstream untargeted metabolomics and prostaglandin E metabolite (PGEM) assays.

Between June 2014 and August 2015, 132 subjects with diagnosed T2D and 35 nondiabetic (ND) controls were enrolled at the University of Wisconsin Hospitals and Clinics (UWHC) to provide biometric parameters, routine clinical labs, and a plasma sample for research purposes. All human subjects research was conducted in accordance with the standards set out by the World Medical Association (WMA) Declaration of Helsinki “Ethical Principles for Medical Research Involving Human Subjects” as approved by the UW Health Sciences IRB (UW 2013-1082). Consent-waived electronic medical record (EMR) search identified potentially-eligible subjects with upcoming morning outpatient appointments in the UWHC Endocrinology Clinic who met baseline inclusion and exclusion criteria. Inclusion criteria were 18-74 years old, not pregnant or lactating, no anemia or grossly abnormal kidney or liver function tests, no known autoimmune diseases or inflammatory disorders, and no diagnosis of diabetes besides T2D. Exclusion criteria were history of transplant, chronic steroid use, or use of COX inhibitors other than prophylactic low-dose aspirin for cardiovascular health more than twice per week during the past 90 days. Patients meeting all inclusion and exclusion criteria were contacted via telephone for verification of EMR data, including dose and frequency of any aspirin therapy. The background and aims of the study were described, including the need for an overnight fast and a laboratory blood draw for research purposes. Patients interested in participating provided initial verbal consent by phone and were instructed to fast for 12 h prior to their upcoming outpatient appointment. If an iatrogenic hypoglycemic event occurred, patients were instructed to treat per standard medical protocol until resolution. The patient’s care provider was contacted to coordinate laboratory test orders, as our study protocol included complete blood count (CBC), complete metabolic panel (CMP), fasting lipid panel, HbA1c, C reactive protein (CRP), and erythrocyte sedimentation rate (ESR), some of which are T2D standard-of-care tests.

Upon presentation at the UWHC lab, a study coordinator described the aims of the study and that participation was voluntary and confirmed eligibility criteria. The study coordinator confirmed the patient had fasted for 12 h. If no greater than 15 grams of carbohydrates were consumed as treatment for any hypoglycemic episode in last 12 hours, the patient remained eligible to provide a blood sample. If greater than 15 grams of carbohydrates were consumed and the patient remained interested in participating, the clinical laboratory appointment was re-scheduled. Consent forms were reviewed and signed by both participants and study coordinators prior to sample procurement and filed in a locked area thereafter. Height, weight, blood pressure, and pulse were measured, and daily omega-3 supplement use was noted. Current T2D medications were confirmed and recorded. A UWHC staff phlebotomist drew blood for clinical lab tests performed as part of the study protocol, followed by a blood draw for research purposes into a 8.5 ml BD™ P800 blood collection tube coated with potassium EDTA and a proprietary mix of protease and esterase inhibitors (BD Biosciences, cat. no. 366421). The research blood tube was gently mixed and immediately refrigerated until transported to the UWHC Central Lab via cold pack, where plasma was separated, aliquoted, and stored at −80°C until used in research lab assays. All subjects providing consent received remuneration in the form of a $25 gift card.

Based on means and SDs from our pre-clinical models, our target recruitment was 137 T2D subjects and 35 non-diabetic (ND) subjects for a power of 80% to see significant differences at alpha = 0.05. One patient carrying a T2D diagnosis prior to their intake appointment being treated with diet/lifestyle modifications had a HgA1c less than 6% and fasting blood glucose value less than 125 mg/dl and was re-assigned to the ND cohort. One patient initially enrolled in the ND group who was being monitored for pre-diabetes had a HbA1c value over 6.5%, and, in coordination with their care provider, informed of their T2D diagnosis and re-assigned to the T2D cohort. Finally, four subjects were excluded from final analysis, all in the T2D group. One subject withdrew consent. One subject did not have clinical labs run. One subject had an elevated white-cell count and, upon follow-up, had been acutely ill on the day of presentation. Lastly, one subject had a low potassium level that triggered the endocrinologist on call to adjudicate. This subject was found to have a history of bariatric surgery and referred to follow-up with their surgeon. Therefore, the final cohort included 132 T2D subjects and 35 ND subjects.

For the current work, only biobanked samples from subjects with well-controlled T2D (HbA1c ≤ 8) under conservative treatment (diet/lifestyle modifications or metformin monotherapy) were desired. 19 of 132 T2D subjects met these criteria, including one subject with a HbA1c of 8.1. Samples from non-diabetic subjects were sorted by BMI, and all samples spanning nearly same BMI range were selected for controls (N=16 of 35).

### Untargeted Plasma Metabolomics Analysis using Flow Injection Ionospray (FIE) Fourier Transform Ion Cyclotron Resonance (FTICR) Mass Spectrometry (MS)

FIE-FTICR MS experiments were performed using a column-free Waters nanoACQUITY UPLC (Waters Corporation, Milford, MA, USA) coupled to a Bruker solariX 12 T FTICR mass spectrometer (Bruker Daltonics, Bremen, Germany) as previously described for mouse plasma samples, with minor modifications^*17, 28*^. Briefly, a 30 *μ*l aliquot of plasma was mixed with 60 *μ*L of chilled liquid chromatography (LC)-MS grade methanol. The samples were then vortexed, mixed with a nutating mixer, and centrifuged. 50 *μ*L of supernatant was mixed with 50 *μ*L of water, and 5 *μ*L of each sample directly injected in triplicate from the Waters nanoACQUITY UPLC into the FTICR MS via 100 *μ*m x 40 cm PEEK tubing. The mobile phase was 50:50 methanol:water with 0.1% formic acid or 10 mM ammonium acetate added for positive or negative modes, respectively, with a flow rate of 20 *μ*l/min. Ions were accumulated for 0.1 s and a 8 M transient size was applied, with 50 scans collected. A m/z range of was set to 40-1500 with 50 m/z Q1 mass. 50 scans were collected for each mass spectrum. Dry gas flow was set to 4 L/min at 150 °C. The largest frequency values for octopole (5 MHz), quadrupole (2 MHz), and transfer hexapole (6 MHz) were used to improve ion transition. Time of flight was set to 0.8 ms. Sweep excitation power was set to 27%. The estimated resolving power at 400 m/z was 190,000. The FTICR MS was calibrated with 1 mM NaTFA in both positive and negative modes before experiments. The mass spectra were processed and analyzed using DataAnalysis 4.3 (Bruker Daltonics, Bremen, Germany) and annotated using the SmartFormula function in MetaboScape 4.0 (Bruker Daltonics, Bremen, Germany) and the METLIN MS/MS database housed by the Scripps Research Institute using a 2 ppm mass error cutoff as previously described^*17, 28*^. The full untargeted metabolomics analysis will be published elsewhere. In this work, the intensity of the peak corresponding with arachidonic acid was compared among groups.

### Prostaglandin E Metabolite (PGEM) Assay

Plasma PGE_2_ levels were quantified using a Prostaglandin E Metabolite (PGEM) ELISA kit (Cayman Chemical Company, cat. no. 514531) according to the manufacturer’s protocol. Briefly, samples were purified by acetone precipitation followed by drying under a gentle nitrogen stream before being re-suspended in an equivalent volume of ELISA buffer. Samples and standards were derivatized overnight and loaded onto 96-well assay plates at a 1:5 dilution in duplicate. No deviations from the ELISA protocol were made, and the optional acidification and ethyl acetate extraction step after derivatization was not necessary.

### Statistical analyses

Statistical analyses were performed with GraphPad Prism v. 9 (GraphPad Software, San Diego, CA) according to the methods described in the figure legends. In all cases, p < 0.05 was considered statistically significant.

## ACKNOWLEDGMENTS

We wish to thank the many present and former members of the Kimple Laboratory who contributed technical assistance or scientific discussion during the course of these experiments, most notably, Jackson Moeller, Renee Buchanan, and Stephanie Blaha. We would also like to thank Dr. Yanlong Zhu, director of the Human Proteomics Program Core Facility, who advised on the technical aspects of this work.

## AUTHOR INFORMATION

**Corresponding Authors**

*Michelle E. Kimple, PhD, Associate Professor of Medicine, Division of Endocrinology, Diabetes, and Metabolism and Faculty Affiliate, Department of Cell and Regenerative Biology, University of Wisconsin School of Medicine and Public Health; Research Health Scientist, William S. Middleton Memorial VA Hospital, 4148 UW Medical Foundation Centennial Building, 1685 Highland Ave., Madison, WI 53705, 608-265-5627

*Dawn B. Davis, MD, PhD, Professor of Medicine and Director of the Comprehensive Diabetes Center, Division of Endocrinology, Diabetes, and Metabolism; Chief of Endocrinology, William S. Middleton Memorial VA Hospital, 4147 UW Medical Foundation Centennial Building, 1685 Highland Ave., Madison, WI 53705, 608-263-2443

## Author Contributions

Conceptualization, EDC, YG, DBD, and MEK; data curation, NAT, RJF, AMW, RN, MD, SP, MP, DBD, and MEK; formal analysis, NAT, RJF, HKS, AMW, CP, BW, and MEK; funding acquisition, RJF, AMW, BW, SP, ALB, EDC, DBD, and MEK; investigation, NAT, RJF, HKS, AMW, BW, SP, AR, JMH, ALB, DCP, RN, MD, MP, and MEK; methodology, NAT, RJF, AMW, BW, YG, DBD, and MEK; project administration, AMW, DBD, and MEK; supervision, NAT, RJF, AMW, YG, DBD, and MEK; visualization, RJF, CP, CEK, and MEK; writing—original draft, RJF, CP, BW, and MEK; writing—review and editing, RJF, CP, MD, CEK, DBD, and MEK. All authors have read and agreed to the published version of the manuscript.

## Supporting Information

The following files are available free of charge.

**Table S1:** Sources and Donor Demographics of islet preparations from Set 1, used for gene expression analyses and insulin secretion assays.

**Table S2:** Complete Correlation analysis of BMI/gene/Insulin content analyses.

**Table S3:** Sources and Donor Demographics of islet preparations from Set 2, used to confirm the relationship of *PTGER3* expression with obesity.

**Table S4:** Quantitative PCR primer sequences

## For Table of Contents use only

Human islet expression levels of Prostaglandin E_2_ synthetic enzymes, but not prostaglandin EP3 receptor, are positively correlated with markers of β-cell function and mass in non-diabetic obesity.

Nathan A. Truchan, Rachel J. Fenske, Harpreet K. Sandhu, Alicia M. Weeks, Chinmai Patibandla, Benjamin Wancewicz, Samantha Pabich, Austin Reuter, Jeffrey M. Harrington, Allison L. Brill, Darby C. Peter, Randall Nall, Michael Daniels, Margaret Punt, Cecilia E. Kaiser, Elizabeth D. Cox, Ying Ge, Dawn B. Davis, and Michelle E. Kimple

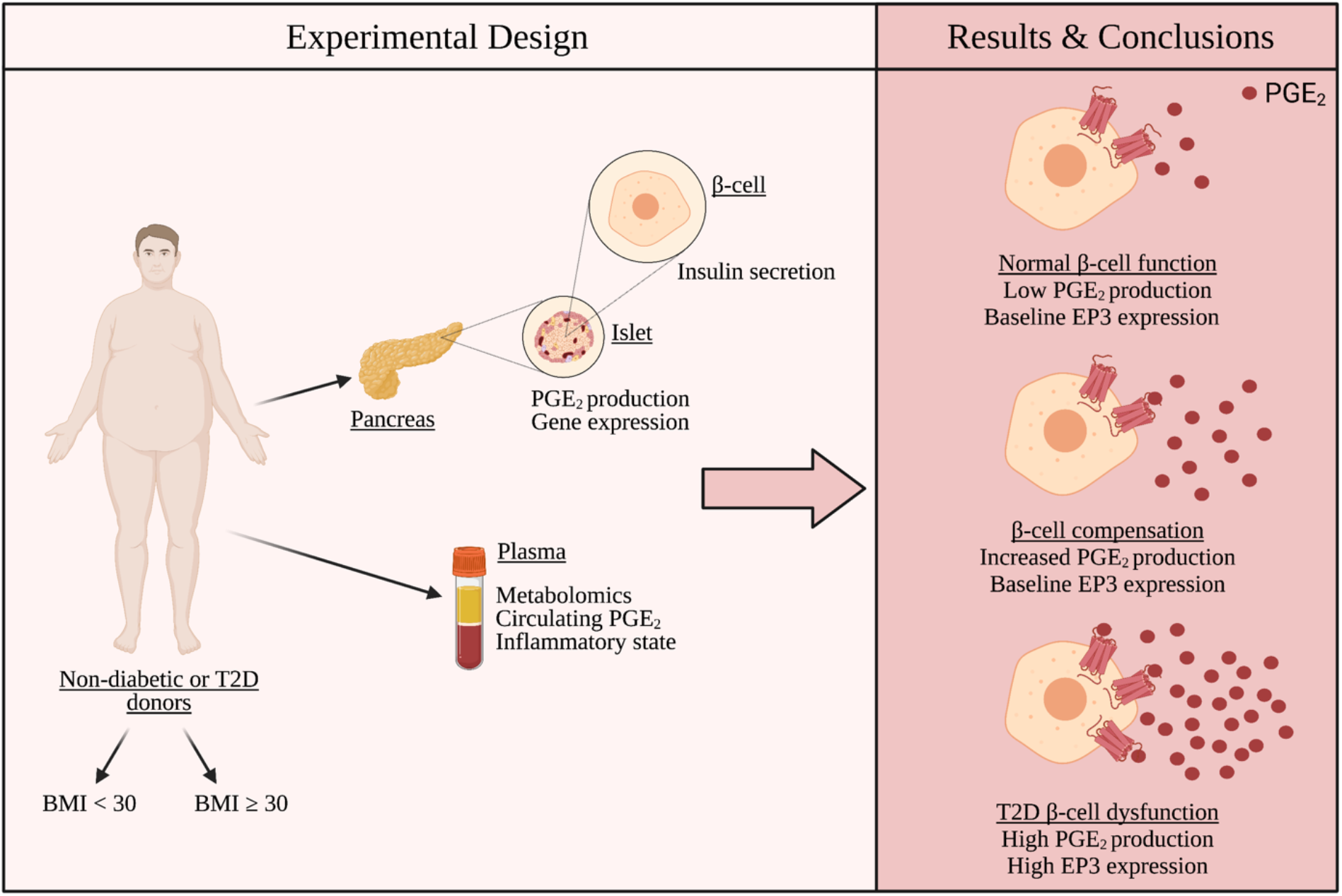

**Description:** Using plasma samples from non-diabetic and type 2 diabetic (T2D) human subjects and human islets isolated from non-diabetic organ donors, we propose step-wise increases in betacell EP3 activity are involved in the progression from beta-cell health to beta-cell compensation and, ultimately, beta-cell failure. Elevated prostaglandin E2 (PGE2) levels may be required for the compensatory response to the inflammation of obesity. Further increases in PGE2 production by hyperglycemia, as well as EP3 expression itself, actively contribute to T2D beta-cell dysfunction. Figure created by Cecilia E. Kaiser with Biorender.com.

## REFERENCES

[1] Jezek, P., Jaburek, M., and Plecita-Hlavata, L. (2019) Contribution of Oxidative Stress and Impaired Biogenesis of Pancreatic beta-Cells to Type 2 Diabetes, Antioxid Redox Signal 31, 722–751.

[2] Alejandro, E. U., Gregg, B., Blandino-Rosano, M., Cras-Meneur, C., and Bernal-Mizrachi, E. (2015) Natural history of beta-cell adaptation and failure in type 2 diabetes, Molecular aspects of medicine 42, 19–41.

[3] Poitout, V., and Robertson, R. P. (2008) Glucolipotoxicity: fuel excess and beta-cell dysfunction, Endocrine reviews 29, 351–366.

[4] Tengholm, A. (2012) Cyclic AMP dynamics in the pancreatic beta-cell, Ups J Med Sci 117, 355–369.

[5] Tengholm, A., and Gylfe, E. (2017) cAMP signalling in insulin and glucagon secretion, Diabetes, obesity & metabolism 19 Suppl 1, 42–53.

[6] Blanchet, E., Van de Velde, S., Matsumura, S., Hao, E., LeLay, J., Kaestner, K., and Montminy, M. (2015) Feedback inhibition of CREB signaling promotes beta cell dysfunction in insulin resistance, Cell reports 10, 1149–1157.

[7] Jhala, U. S., Canettieri, G., Screaton, R. A., Kulkarni, R. N., Krajewski, S., Reed, J., Walker, J., Lin, X., White, M., and Montminy, M. (2003) cAMP promotes pancreatic beta-cell survival via CREB-mediated induction of IRS2, Genes & development 17, 1575–1580.

[8] Dyachok, O., Idevall-Hagren, O., Sagetorp, J., Tian, G., Wuttke, A., Arrieumerlou, C., Akusjarvi, G., Gylfe, E., and Tengholm, A. (2008) Glucose-induced cyclic AMP oscillations regulate pulsatile insulin secretion, Cell metabolism 8, 26–37.

[9] Carboneau, B. A., Breyer, R. M., and Gannon, M. (2017) Regulation of pancreatic beta-cell function and mass dynamics by prostaglandin signaling, J Cell Commun Signal 11, 105–116.

[10] Schaid, M. D., Green, C. L., Peter, D. C., Gallagher, S. J., Guthery, E., Carbajal, K. A., Harrington, J. M., Kelly, G. M., Reuter, A., Wehner, M. L., Brill, A. L., Neuman, J. C., Lamming, D. W., and Kimple, M. E. (2020) Agonist-independent Galphaz activity negatively regulates beta-cell compensation in a diet-induced obesity model of type 2 diabetes, J Biol Chem.

[11] Black, M. A., Heick, H. M., and Begin-Heick, N. (1988) Abnormal regulation of cAMP accumulation in pancreatic islets of obese mice, The American journal of physiology 255,E833-838.

[12] Portela-Gomes, G. M., and Abdel-Halim, S. M. (2002) Overexpression of Gs proteins and adenylyl cyclase in normal and diabetic islets, Pancreas 25, 176–181.

[13] Kimple, M. E., Keller, M. P., Rabaglia, M. R., Pasker, R. L., Neuman, J. C., Truchan, N. A., Brar, H. K., and Attie, A. D. (2013) Prostaglandin E2 receptor, EP3, is induced in diabetic islets and negatively regulates glucose- and hormone-stimulated insulin secretion, Diabetes 62, 1904–1912.

[14] Carboneau, B. A., Allan, J. A., Townsend, S. E., Kimple, M. E., Breyer, R. M., and Gannon, M. (2017) Opposing effects of prostaglandin E2 receptors EP3 and EP4 on mouse and human beta-cell survival and proliferation, Molecular metabolism 6, 548–559.

[15] Amior, L., Srivastava, R., Nano, R., Bertuzzi, F., and Melloul, D. (2019) The role of Cox-2 and prostaglandin E2 receptor EP3 in pancreatic beta-cell death, FASEB J 33, 4975–4986.

[16] Kimple, M. E., Moss, J. B., Brar, H. K., Rosa, T. C., Truchan, N. A., Pasker, R. L., Newgard, C. B., and Casey, P. J. (2012) Deletion of GalphaZ protein protects against diet-induced glucose intolerance via expansion of beta-cell mass, The Journal of biological chemistry 287, 20344–20355.

[17] Schaid, M. D., Zhu, Y., Richardson, N. E., Patibandla, C., Ong, I. M., Fenske, R. J., Neuman, J. C., Guthery, E., Reuter, A., Sandhu, H. K., Fuller, M. H., Cox, E. D., Davis, D. B., Layden, B. T., Brasier, A. R., Lamming, D. W., Ge, Y., and Kimple, M. E. (2021) Systemic Metabolic Alterations Correlate with Islet-Level Prostaglandin E2 Production and Signaling Mechanisms That Predict beta-Cell Dysfunction in a Mouse Model of Type 2 Diabetes, Metabolites 11.

[18] Neuman, J. C., Schaid, M. D., Brill, A. L., Fenske, R. J., Kibbe, C. R., Fontaine, D. A., Sdao, S. M., Brar, H. K., Connors, K. M., Wienkes, H. N., Eliceiri, K. W., Merrins, M. J., Davis, D. B., and Kimple, M. E. (2017) Enriching Islet Phospholipids With Eicosapentaenoic Acid Reduces Prostaglandin E2 Signaling and Enhances Diabetic beta-Cell Function, Diabetes 66, 1572–1585.

[19] Sjoholm, A. (1996) Prostaglandins inhibit pancreatic beta-cell replication and long-term insulin secretion by pertussis toxin-insensitive mechanisms but do not mediate the actions of interleukin-1 beta, Biochimica et biophysica acta 1313, 106–110.

[20] Tran, P. O., Gleason, C. E., Poitout, V., and Robertson, R. P. (1999) Prostaglandin E(2) mediates inhibition of insulin secretion by interleukin-1beta, The Journal of biological chemistry 274, 31245–31248.

[21] Tran, P. O., Gleason, C. E., and Robertson, R. P. (2002) Inhibition of interleukin-1beta-induced COX-2 and EP3 gene expression by sodium salicylate enhances pancreatic islet beta-cell function, Diabetes 51, 1772–1778.

[22] Abdel-Magid, A. F. (2015) Prostaglandin EP3 Receptor Antagonists May Provide Novel Treatment for Diabetes, ACS medicinal chemistry letters 6, 626–627.

[23] Robertson, R. P. (2017) The COX-2/PGE2/EP3/Gi/o/cAMP/GSIS Pathway in the Islet: The Beat Goes On, Diabetes 66, 1464–1466.

[24] Schaid, M. D., Wisinski, J. A., and Kimple, M. E. (2017) The EP3 Receptor/Gz Signaling Axis as a Therapeutic Target for Diabetes and Cardiovascular Disease, The AAPS journal 19, 1276–1283.

[25] Kimple, M. E., Neuman, J. C., Linnemann, A. K., and Casey, P. J. (2014) Inhibitory G proteins and their receptors: emerging therapeutic targets for obesity and diabetes, Exp Mol Med 46, e102.

[26] Persaud, S. J., Burns, C. J., Belin, V. D., and Jones, P. M. (2004) Glucose-induced regulation of COX-2 expression in human islets of Langerhans, Diabetes 53 Suppl 1,S190-192.

[27] Sandhu, H. K., Neuman, J. C., Schaid, M. D., Davis, S. E., Connors, K. M., Challa, R., Guthery, E., Fenske, R. J., Patibandla, C., Breyer, R. M., and Kimple, M. E. (2021) Rat prostaglandin EP3 receptor is highly promiscuous and is the sole prostanoid receptor family member that regulates INS-1 (832/3) cell glucose-stimulated insulin secretion, Pharmacol Res Perspect 9, e00736.

[28] Zhu, Y., Wancewicz, B., Schaid, M., Tiambeng, T. N., Wenger, K., Jin, Y., Heyman, H., Thompson, C. J., Barsch, A., Cox, E. D., Davis, D. B., Brasier, A. R., Kimple, M. E., and Ge, Y. (2021) Ultrahigh-Resolution Mass Spectrometry-Based Platform for Plasma Metabolomics Applied to Type 2 Diabetes Research, J Proteome Res 20, 463–473.

[29] Heitmeier, M. R., Kelly, C. B., Ensor, N. J., Gibson, K. A., Mullis, K. G., Corbett, J. A., and Maziasz, T. J. (2004) Role of cyclooxygenase-2 in cytokine-induced beta-cell dysfunction and damage by isolated rat and human islets, The Journal of biological chemistry 279, 53145–53151.

[30] Kwon, G., Corbett, J. A., Hauser, S., Hill, J. R., Turk, J., and McDaniel, M. L. (1998) Evidence for involvement of the proteasome complex (26S) and NFkappaB in IL-1beta-induced nitric oxide and prostaglandin production by rat islets and RINm5F cells, Diabetes 47, 583–591.

[31] Corbett, J. A., Kwon, G., Turk, J., and McDaniel, M. L. (1993) IL-1 beta induces the coexpression of both nitric oxide synthase and cyclooxygenase by islets of Langerhans: activation of cyclooxygenase by nitric oxide, Biochemistry 32, 13767–13770.

[32] Neuman, J. C., Fenske, R. J., and Kimple, M. E. (2017) Dietary polyunsaturated fatty acids and their metabolites: Implications for diabetes pathophysiology, prevention, and treatment, Nutrition and healthy aging 4, 127–140.

[33] Brill, A. L., Wisinski, J. A., Cadena, M. T., Thompson, M. F., Fenske, R. J., Brar, H. K., Schaid, M. D., Pasker, R. L., and Kimple, M. E. (2016) Synergy Between Galphaz Deficiency and GLP-1 Analog Treatment in Preserving Functional beta-Cell Mass in Experimental Diabetes, Molecular endocrinology 30, 543–556.

[34] Bastard, J. P., Maachi, M., Lagathu, C., Kim, M. J., Caron, M., Vidal, H., Capeau, J., and Feve, B. (2006) Recent advances in the relationship between obesity, inflammation, and insulin resistance, Eur Cytokine Netw 17, 4–12.

[35] Eder, K., Baffy, N., Falus, A., and Fulop, A. K. (2009) The major inflammatory mediator interleukin-6 and obesity, Inflamm Res 58, 727–736.

[36] Han, M. S., White, A., Perry, R. J., Camporez, J. P., Hidalgo, J., Shulman, G. I., and Davis, R. J. (2020) Regulation of adipose tissue inflammation by interleukin 6, Proc Natl Acad Sci USA 117, 2751–2760.

[37] Truchan, N. A., Brar, H. K., Gallagher, S. J., Neuman, J. C., and Kimple, M. E. (2015) A single-islet microplate assay to measure mouse and human islet insulin secretion, Islets 7, e1076607.

[38] Wang, J., Yang, X., and Zhang, J. (2016) Bridges between mitochondrial oxidative stress, ER stress and mTOR signaling in pancreatic beta cells, Cellular signalling 28, 1099–1104.

[39] Tabatabaie, T., Vasquez-Weldon, A., Moore, D. R., and Kotake, Y. (2003) Free radicals and the pathogenesis of type 1 diabetes: beta-cell cytokine-mediated free radical generation via cyclooxygenase-2, Diabetes 52, 1994–1999.

[40] Ceddia, R. P., Lee, D., Maulis, M. F., Carboneau, B. A., Threadgill, D. W., Poffenberger, G., Milne, G., Boyd, K. L., Powers, A. C., McGuinness, O. P., Gannon, M., and Breyer, R. M. (2016) The PGE2 EP3 Receptor Regulates Diet-Induced Adiposity in Male Mice, Endocrinology 157, 220–232.

[41] McRae, J. R., Day, R. P., Metz, S. A., Halter, J. B., Ensinck, J. W., and Robertson, R. P. (1985) Prostaglandin E2 metabolite levels during diabetic ketoacidosis, Diabetes 34, 761–766.

[42] Elisia, I., Lam, V., Cho, B., Hay, M., Li, M. Y., Kapeluto, J., Elliott, T., Harris, D., Bu, L., Jia, W., Leung, H., Mohn, W., and Krystal, G. (2020) Exploratory examination of inflammation state, immune response and blood cell composition in a human obese cohort to identify potential markers predicting cancer risk, PLoS One 15, e0228633.

[43] Tans, R., Bande, R., van Rooij, A., Molloy, B. J., Stienstra, R., Tack, C. J., Wevers, R. A., Wessels, H., Gloerich, J., and van Gool, A. J. (2020) Evaluation of cyclooxygenase oxylipins as potential biomarker for obesity-associated adipose tissue inflammation and type 2 diabetes using targeted multiple reaction monitoring mass spectrometry, Prostaglandins Leukot Essent Fatty Acids 160, 102157.

[44] Xia, F., He, C., Ren, M., Xu, F. G., and Wan, J. B. (2020) Quantitative profiling of eicosanoids derived from n-6 and n-3 polyunsaturated fatty acids by twin derivatization strategy combined with LC-MS/MS in patients with type 2 diabetes mellitus, Anal Chim Acta 1120, 24–35.

[45] Pawelzik, S. C., Avignon, A., Idborg, H., Boegner, C., Stanke-Labesque, F., Jakobsson, P. J., Sultan, A., and Back, M. (2019) Urinary prostaglandin D2 and E2 metabolites associate with abdominal obesity, glucose metabolism, and triglycerides in obese subjects, Prostaglandins Other Lipid Mediat 145, 106361.

[46] Fenske, R. J., Cadena, M. T., Harenda, Q. E., Wienkes, H. N., Carbajal, K., Schaid, M. D., Laundre, E., Brill, A. L., Truchan, N. A., Brar, H., Wisinski, J., Cai, J., Graham, T. E., Engin, F., and Kimple, M. E. (2017) The Inhibitory G Protein alpha-Subunit, Galphaz, Promotes Type 1 Diabetes-Like Pathophysiology in NOD Mice, Endocrinology 158, 1645–1658.

[47] Parazzoli, S., Harmon, J. S., Vallerie, S. N., Zhang, T., Zhou, H., and Robertson, R. P. (2012) Cyclooxygenase-2, not microsomal prostaglandin E synthase-1, is the mechanism for interleukin-1beta-induced prostaglandin E2 production and inhibition of insulin secretion in pancreatic islets, J Biol Chem 287, 32246–32253.

[48] Neuman, J. C., and Kimple, M. E. (2013) The EP3 Receptor: Exploring a New Target for Type 2 Diabetes Therapeutics, J Endocrinol Diabetes Obes 1.

[49] Robertson, R. P. (1998) Dominance of cyclooxygenase-2 in the regulation of pancreatic islet prostaglandin synthesis, Diabetes 47, 1379–1383.

[50] Fenske, R. J., and Kimple, M. E. (2018) Targeting dysfunctional beta-cell signaling for the potential treatment of type 1 diabetes mellitus, Experimental biology and medicine 243, 586–591.

[51] Choi, S. E., Choi, K. M., Yoon, I. H., Shin, J. Y., Kim, J. S., Park, W. Y., Han, D. J., Kim, S. C., Ahn, C., Kim, J. Y., Hwang, E. S., Cha, C. Y., Szot, G. L., Yoon, K. H., and Park, C. G. (2004) IL-6 protects pancreatic islet beta cells from pro-inflammatory cytokines-induced cell death and functional impairment in vitro and in vivo, Transpl Immunol 13, 43–53.

[52] da Silva Krause, M., Bittencourt, A., Homem de Bittencourt, P. I., Jr., McClenaghan, N. H., Flatt, P. R., Murphy, C., and Newsholme, P. (2012) Physiological concentrations of interleukin-6 directly promote insulin secretion, signal transduction, nitric oxide release, and redox status in a clonal pancreatic beta-cell line and mouse islets, J Endocrinol 214, 301–311.

[53] Linnemann, A. K., Blumer, J., Marasco, M. R., Battiola, T. J., Umhoefer, H. M., Han, J. Y., Lamming, D. W., and Davis, D. B. (2017) Interleukin 6 protects pancreatic beta cells from apoptosis by stimulation of autophagy, FASEB journal: official publication of the Federation of American Societies for Experimental Biology 31, 4140–4152.

[54] Marasco, M. R., Conteh, A. M., Reissaus, C. A., Cupit, J. E. t., Appleman, E. M., Mirmira, R. G., and Linnemann, A. K. (2018) Interleukin-6 Reduces beta-Cell Oxidative Stress by Linking Autophagy With the Antioxidant Response, Diabetes 67, 1576–1588.

[55] Ellingsgaard, H., Ehses, J. A., Hammar, E. B., Van Lommel, L., Quintens, R., Martens, G., Kerr-Conte, J., Pattou, F., Berney, T., Pipeleers, D., Halban, P. A., Schuit, F. C., and Donath, M. Y. (2008) Interleukin-6 regulates pancreatic alpha-cell mass expansion, Proceedings of the National Academy of Sciences of the United States of America 105, 13163–13168.

[56] Bagga, D., Wang, L., Farias-Eisner, R., Glaspy, J. A., and Reddy, S. T. (2003) Differential effects of prostaglandin derived from omega-6 and omega-3 polyunsaturated fatty acids on COX-2 expression and IL-6 secretion, Proceedings of the National Academy of Sciences of the United States of America 100, 1751–1756.

[57] Sorli, C. H., Zhang, H. J., Armstrong, M. B., Rajotte, R. V., Maclouf, J., and Robertson, R. P. (1998) Basal expression of cyclooxygenase-2 and nuclear factor-interleukin 6 are dominant and coordinately regulated by interleukin 1 in the pancreatic islet, Proceedings of the National Academy of Sciences of the United States of America 95, 1788–1793.

[58] Henquin, J. C. (2018) Influence of organ donor attributes and preparation characteristics on the dynamics of insulin secretion in isolated human islets, Physiol Rep 6.

[59] Brandhorst, H., Brandhorst, D., Hering, B. J., Federlin, K., and Bretzel, R. G. (1995) Body mass index of pancreatic donors: a decisive factor for human islet isolation, Exp Clin Endocrinol Diabetes 103 Suppl 2, 23–26.

