## Supplementary Table for "Human islet expression levels of Prostaglandin E_2_ synthetic enzymes, but not prostaglandin EP3 receptor, are positively correlated with markers of β-cell function and mass in non-diabetic obesity"

| Table S1: Primer sequences used for quantitative gene expression analyses. | | | |
|---|---|---|---|
| **Protein** | **Gene Symbol** | **Primer Sequences** | **Species Selectivity** |
| **β-actin** | *ACTB* | F: ACCACACCTTCTACAATGAGC<br>R: GATAGCACAGCCTGGATAGC | human |
| **Interleukin-6**<br>**(IL-6)** | *IL6* | F: GTGCTCTTGGTGAGGAAGTT<br>R: TTCTGGGACTCCTGGGAATA | human |
| **Cyclooxygenase 1**<br>**(COX-1) (aka PTGS1)** | *PTGS1* | F: GTTCAACACCTCCATGTTGG<br>R: CCACAGCCACATGCAGGATG | human |
| **Cyclooxygenase 2**<br>**(COX-2) (aka PTGS2)** | *PTGS2* | F: CGAGGTGTATGTATGAGTGTGG<br>R: CAAAAATTCCGGTGTTGAGCAG | human |
| **Prostaglandin E synthase**<br>**(PTGES)** | *PTGES* | F: TGGTCATCAAGATGTACGTGGTGGC<br>R: TAGATGGTCTCCATGTCGTTCCGGT | human |
| **Prostaglandin E synthase 2**<br>**(PTGES2)** | *PTGES2* | F: ACCTCTATGAGGCTGCTGACAAGT<br>R: CATACACCGCCAAATCAGCGAGAT | human |
| **Prostaglandin E synthase 3**<br>**(Ptges3)** | *PTGES3* | F: AAAGGAGAATCTGGCCAGTCATGG<br>R: TCCTCATCACCACCCATGTTGTTC | human |
| **Prostaglandin EP3 receptor**<br>**(EP3)** | *PTGER3* | F: TCACCTTTTCCTGCAACCTG<br>R: ACGCACATGATCCCCATAAG | human |

Supplementary Table 2: Islet Data for Preparations in Set 2 (Received Mar 2013-May 2015)

| Unique Identifier | AGE | SEX | BMI | HbA1c | Origin | Islet Isolation Center | Donor History of diabetes? | Assay |
|---|---|---|---|---|---|---|---|---|
| SAMN08930526 | 42 | M | 22.8 | Not Reported | IIDP | University of Illinois | No | Gene Expression and GSIS |
| SAMN08775090 | 32 | M | 23.1 | Not Reported | IIDP | The Scharp-Lacy Research Institute | No | Gene Expression |
| SAMN08784299 | 23 | M | 23.9 | Not Reported | IIDP | The Scharp-Lacy Research Institute | No | Gene Expression and GSIS |
| SAMN08930530 | 48 | F | 24.2 | Not Reported | IIDP | University of Illinois | No | Gene Expression |
| SAMN08930578 | 51 | M | 24.7 | Not Reported | IIDP | University of Illinois | No | Gene Expression |
| SAMN08784380 | 50 | M | 25.7 | Not Reported | IIDP | University of Miami | No | Gene Expression and GSIS |
| SAMN08774963 | 39 | F | 25.9 | Not Reported | IIDP | The Scharp-Lacy Research Institute | No | Gene Expression and GSIS |
| SAMN08775094 | 36 | M | 26 | Not Reported | IIDP | The Scharp-Lacy Research Institute | No | Gene Expression and GSIS |
| SAMN08784374 | 24 | F | 26.6 | Not Reported | IIDP | University of Miami | No | Gene Expression |
| SAMN08784309 | 25 | M | 26.6 | Not Reported | IIDP | The Scharp-Lacy Research Institute | No | Gene Expression and GSIS |
| SAMN08783919 | 19 | M | 26.9 | Not Reported | IIDP | University of Wisconsin | No | Gene Expression and GSIS |
| SAMN08783909 | 53 | M | 27.2 | Not Reported | IIDP | The Scharp-Lacy Research Institute | No | Gene Expression and GSIS |
| SAMN08498443 | 61 | F | 27.6 | Not Reported | IIDP | Southern California Islet Cell Resource Center | No | Gene Expression |
| SAMN08783908 | 52 | M | 29.1 | Not Reported | IIDP | The Scharp-Lacy Research Institute | No | Gene Expression and GSIS |
| SAMN08776514 | 48 | F | 29.2 | Not Reported | IIDP | University of Wisconsin | No | Gene Expression and GSIS |
| SAMN08776501 | 46 | M | 29.3 | Not Reported | IIDP | The Scharp-Lacy Research Institute | No | Gene Expression and GSIS |
| SAMN08784318 | 43 | M | 29.6 | Not Reported | IIDP | University of Pennsylvania | No | Gene Expression |
| SAMN08775091 | 38 | F | 30.6 | Not Reported | IIDP | The Scharp-Lacy Research Institute | No | Gene Expression |
| SAMN08784301 | 43 | M | 30.6 | Not Reported | IIDP | The Scharp-Lacy Research Institute | No | Gene Expression and GSIS |
| SAMN08774955 | 48 | M | 30.7 | Not Reported | IIDP | Southern California Islet Cell Resource Center | No | Gene Expression |
| SAMN08776527 | 58 | F | 31.1 | Not Reported | IIDP | University of Wisconsin | No | Gene Expression |
| SAMN08774969 | 59 | F | 31.3 | Not Reported | IIDP | The Scharp-Lacy Research Institute | No | Gene Expression and GSIS |
| SAMN08774961 | 52 | F | 31.4 | Not Reported | IIDP | Southern California Islet Cell Resource Center | No | Gene Expression and GSIS |
| SAMN08930577 | 50 | M | 31.7 | Not Reported | IIDP | University of Illinois | No | Gene Expression |
| SAMN08774814 | 45 | F | 32.9 | Not Reported | IIDP | University of Wisconsin | No | Gene Expression |
| SAMN08774197 | 52 | M | 33.3 | 4.6 | IIDP | University of Wisconsin | No | Gene Expression |
| SAMN08776521 | 36 | M | 33.8 | Not Reported | IIDP | The Scharp-Lacy Research Institute | No | Gene Expression |
| SAMN08784305 | 52 | M | 34.3 | Not Reported | IIDP | The Scharp-Lacy Research Institute | No | Gene Expression and GSIS |
| SAMN08784314 | 36 | F | 34.8 | Not Reported | IIDP | The Scharp-Lacy Research Institute | No | Gene Expression and GSIS |
| SAMN08775085 | 58 | M | 34.8 | Not Reported | IIDP | University of Wisconsin | No | Gene Expression and GSIS |
| SAMN08776506 | 40 | M | 35.4 | Not Reported | IIDP | University of Pennsylvania | No | Gene Expression and GSIS |
| SAMN08783899 | 25 | M | 35.7 | Not Reported | IIDP | University of Wisconsin | No | Gene Expression and GSIS |
| SAMN08775087 | 45 | M | 36.4 | Not Reported | IIDP | University of Wisconsin | No | Gene Expression |
| SAMN08784317 | 21 | M | 37 | Not Reported | IIDP | University of Wisconsin | No | Gene Expression and GSIS |
| SAMN08774896 | 63 | M | 38.6 | Not Reported | IIDP | University of Wisconsin | No | Gene Expression |
| SAMN08783906 | 40 | M | 38.9 | Not Reported | IIDP | University of Wisconsin | No | Gene Expression and GSIS |
| SAMN08783912 | 51 | M | 38.9 | Not Reported | IIDP | University of Wisconsin | No | Gene Expression |
| SAMN08775048 | 32 | F | 39.4 | Not Reported | IIDP | University of Wisconsin | No | Gene Expression and GSIS |
| SAMN08775030 | 30 | M | 43.7 | Not Reported | IIDP | University of Wisconsin | No | Gene Expression |
| SAMN08774465 | 38 | F | 44.7 | 4.6 | IIDP | University of Wisconsin | No | Gene Expression |

**Statistical Results of Gene vs. BMI Analyses - Linear Regression**

| IL6 | PTGS1 | PTGS2 | PTGES | PTGES2 | PTGES3 | PTGER3 | Linear Fit |
|---|---|---|---|---|---|---|---|
| 0.2023 | 0.06305 | 0.1396 | 0.04946 | -0.01448 | -0.01404 | 0.02448 | **Slope** |
| 0.0875 | 0.06423 | 0.06778 | 0.07391 | 0.04044 | 0.03253 | 0.04567 | **Std. Error** |
| 0.02514 to 0.3794 | -0.06710 to 0.1932 | 0.002106 to 0.2770 | -0.1002 to 0.1991 | -0.09635 to 0.06739 | -0.07995 to 0.05188 | -0.06798 to 0.1169 | **95% CI** |
| 0.1233 | 0.02538 | 0.1054 | 0.01165 | 0.003361 | 0.005007 | 0.007502 | **R-square** |
| 0.0263 | 0.3327 | 0.0468 | 0.5074 | 0.7223 | 0.6686 | 0.5951 | **P-value** |

**Statistical Results of Gene vs. BMI Analyses - Obesity Status**

| IL6 | PTGS1 | PTGS2 | PTGES | PTGES2 | PTGES3 | PTGER3 | Obesity Status |
|---|---|---|---|---|---|---|---|
| -12.28 | -11.2 | -7.11 | -9.396 | -6.219 | -3.686 | -9.332 | **BMI < 30** |
| -9.795 | -10.79 | -5.509 | -9.438 | -6.27 | -3.613 | -9.163 | **BMI ≥ 30** |
| 2.487 ± 0.9526 | 0.4075 ± 0.7242 | 1.601 ± 0.7524 | -0.04194 ± 0.8229 | -0.05118 ± 0.4483 | 0.07374 ± 0.3642 | 0.1683 ± 0.5067 | **Difference ± SEM** |
| 0.5582 to 4.415 | -1.060 to 1.875 | 0.07539 to 3.127 | -1.708 to 1.624 | -0.9588 to 0.8564 | -0.6641 to 0.8116 | -0.8575 to 1.194 | **95% CI** |
| 0.0129 | 0.577 | 0.0402 | 0.9596 | 0.9097 | 0.8406 | 0.7416 | **P-value** |

| Supplementary Table 4: Islet Data for Preparations in Set 2 (Received Oct 2010-Feb 2012) | | | | | | | | |
|---|---|---|---|---|---|---|---|---|
| **Unique Identifier** | **AGE** | **SEX** | **BMI** | **HbA1c** | **Origin** | **Islet Isolation Center** | **Donor History of diabetes?** | **Assay** |
| SAMN08786321 | 27 | M | 19 | Not Reported | IIDP | The Scharp-Lacy Research Institute | No | Gene Expression |
| SAMN08933943 | 55 | M | 19.2 | Not Reported | IIDP | Emory University | No | Gene Expression |
| Not Available* | 58 | F | 20.1 | Not Reported | Beta-Pro | Beta-Pro | No | Gene Expression |
| SAMN08933951 | 51 | F | 23.1 | Not Reported | IIDP | Massachusetts General Hospital | No | Gene Expression |
| SAMN08930052 | 61 | F | 23.2 | Not Reported | IIDP | University of Illinois | No | Gene Expression |
| SAMN08786253 | 65 | M | 24.5 | Not Reported | IIDP | The Scharp-Lacy Research Institute | No | Gene Expression |
| SAMN08933947 | 44 | M | 24.7 | Not Reported | IIDP | Massachusetts General Hospital | No | Gene Expression |
| SAMN08786220 | 48 | M | 24.7 | Not Reported | IIDP | Southern California Islet Cell Resource Center | No | Gene Expression |
| SAMN08933959 | 32 | M | 25.9 | Not Reported | IIDP | University of Pittsburgh | No | Gene Expression |
| SAMN08933959 | 32 | M | 25.9 | Not Reported | IIDP | University of Pittsburgh | No | Gene Expression |
| SAMN08933939 | 40 | F | 26 | Not Reported | IIDP | Massachusetts General Hospital | No | Gene Expression |
| SAMN08930051 | 44 | M | 26.3 | Not Reported | IIDP | University of Illinois | No | Gene Expression |
| SAMN08930051 | 44 | M | 26.3 | Not Reported | IIDP | University of Illinois | No | Gene Expression |
| SAMN08933945 | 39 | M | 27.4 | Not Reported | IIDP | Massachusetts General Hospital | No | Gene Expression |
| SAMN08786303 | 20 | M | 28.3 | Not Reported | IIDP | The Scharp-Lacy Research Institute | No | Gene Expression |
| Not Available* | 44 | F | 29.7 | Not Reported | Beta-Pro | Beta-Pro | No | Gene Expression |
| SAMN08786270 | 38 | M | 29.8 | Not Reported | IIDP | The Scharp-Lacy Research Institute | No | Gene Expression |
| SAMN08786342 | 64 | F | 30 | Not Reported | IIDP | The Scharp-Lacy Research Institute | No | Gene Expression |
| SAMN08786201 | 27 | M | 30 | Not Reported | IIDP | University of Miami | No | Gene Expression |
| SAMN08933935 | 29 | M | 30.2 | Not Reported | IIDP | Massachusetts General Hospital | No | Gene Expression |
| SAMN08933935 | 29 | M | 30.2 | Not Reported | IIDP | Massachusetts General Hospital | No | Gene Expression |
| SAMN08785758 | 56 | M | 30.9 | Not Reported | IIDP | The Scharp-Lacy Research Institute | No | Gene Expression |
| SAMN08786288 | 20 | M | 31.3 | Not Reported | IIDP | University of Miami | No | Gene Expression |
| SAMN08786302 | 51 | F | 31.6 | Not Reported | IIDP | University of Pennsylvania | No | Gene Expression |
| SAMN08786302 | 51 | F | 31.6 | Not Reported | IIDP | University of Pennsylvania | No | Gene Expression |
| SAMN08786218 | 17 | M | 32.5 | Not Reported | IIDP | Southern California Islet Cell Resource Center | No | Gene Expression |
| SAMN08933824 | 49 | F | 33.1 | Not Reported | IIDP | University of Minnesota | No | Gene Expression |
| SAMN08786257 | 48 | F | 33.1 | Not Reported | IIDP | University of Miami | No | Gene Expression |
| SAMN08933825 | 48 | M | 34.4 | Not Reported | IIDP | University of Minnesota | No | Gene Expression |
| Not Available* | 37 | F | 34.4 | Not Reported | Beta-Pro | Beta-Pro | No | Gene Expression |
| SAMN08786315 | 28 | M | 34.5 | Not Reported | IIDP | University of Pennsylvania | No | Gene Expression |
| SAMN08786267 | 37 | F | 34.5 | Not Reported | IIDP | Southern California Islet Cell Resource Center | No | Gene Expression |
| SAMN08933940 | 64 | M | 34.8 | Not Reported | IIDP | Massachusetts General Hospital | No | Gene Expression |
| SAMN08933869 | 16 | F | 35.1 | Not Reported | IIDP | Emory University | No | Gene Expression |
| SAMN08786278 | 54 | F | 36 | Not Reported | IIDP | Southern California Islet Cell Resource Center | No | Gene Expression |
| SAMN08786271 | 40 | F | 36.4 | Not Reported | IIDP | The Scharp-Lacy Research Institute | No | Gene Expression |
| SAMN08930084 | 42 | M | 41.3 | Not Reported | IIDP | University of Illinois | No | Gene Expression |
| SAMN08930084 | 42 | M | 41.3 | Not Reported | IIDP | University of Illinois | No | Gene Expression |
| SAMN08786143 | 29 | M | 42 | Not Reported | IIDP | The Scharp-Lacy Research Institute | No | Gene Expression |
| SAMN08786252 | 36 | F | 42.2 | Not Reported | IIDP | Southern California Islet Cell Resource Center | No | Gene Expression |

*Beta-pro LLC is no longer in operation and Unique Identifiers cannot be obtained
